# Do Symptom Domains Have Similar Cellular Underpinnings Across Psychiatric Diagnoses: Evidence from 3D Hippocampal MR Spectroscopy

**DOI:** 10.64898/2026.04.27.721016

**Authors:** Eugene Ruby, Oded Gonen, Eyal Lotan, Assaf Tal, Henry Rusinek, Jose C Clemente Litran, Jessica Robinson-Papp, Katherine H. Karlsgodt, Dolores Malaspina

## Abstract

**Introduction:** The NIMH Research Domain Criteria (RDoC) posits similar cellular pathologies for particular symptom domains across diagnostic categories. Conversely, knowledge that these differ could advance treatment discovery, especially for affective and non-affective psychoses, as studies usually intermix them.

**Methods:** We tested this by relating metabolite biomarker concentrations for cellular pathologies from whole hippocampal proton magnetic resonance spectroscopic imaging (^1^H MRSI) to symptom levels from the original and five factor PANSS, and the Hamilton Depression and Young Mania Scales. Biomarkers included: NAA (neuronal integrity); Cr (energy metabolism); Cho (membrane turnover); mI (glia); and Glx (glutaminergic neurotransmission). Participants were 26 healthy controls; 22 non-psychotic affective cases (NP-aff); and 33 with psychosis, including 20 schizophrenia (Scz) and 13 affective psychosis (aff-P) cases.

**Results:** PANSS activation factor was related to reductions in all cellular component biomarkers in Scz, including glia (ρ=−.84, p<.001), membrane turnover (ρ=−.45, p=.047), neural integrity (ρ=−.54, p=.014), glutaminergic neurotransmission (ρ=−.66, p=.037), and energy metabolism (ρ=−.50, p=.024), but only to energy metabolism in NP-aff (ρ=−.46, p=.030). Biomarkers for mood symptoms also varied across categories, suggesting gliosis for mania (ρ=.49, p=.025) and depression in HC (ρ=.55, p=.012), but increased membrane turnover for mania in aff-P (ρ=.65, p=.015), and decreased neural integrity (ρ=−.45, p=.047)and energy metabolism (ρ=−.45, p=.0496) for depression in Scz. In contrast, negative symptoms and autistic preoccupation were related to reduced glia in both NP-aff (ρ=−.67, p=.005; ρ=−.55, p=.029, respectively) and aff-P (ρ=−.77, p=.042; ρ=−.78, p=.040, respectively). Autistic preoccupation in Scz was related to both reduced glia (ρ=−.55, p=.044)and membrane turnover (ρ=−.45, p=.046). Only Scz showed a significant finding for positive symptoms; reduced membrane turnover (ρ=−.52, p=.018).

**Discussion:** These results suggest both distinct and similar cellular pathologies for symptoms across diagnoses, including affective and non-affective psychoses. The differences support categorizing disorders and stratifying different psychoses in research rather than transdiagnostic approaches.

## INTRODUCTION

A major challenge in clinical psychiatry research is the possibility that similar symptoms arise from distinct cellular pathologies across different diagnostic categories. This possibility is in contrast with the NIMH Research Domain Criteria (RDoC) framework, which posits that common cellular pathologies underlie symptom domains across psychiatric categories [1]. The assumption of shared pathology is especially relevant for studies of psychotic disorders, which typically group different psychoses together (e.g., schizophrenia and affective psychoses). This transdiagnostic approach of intermixing different psychoses derives from the nineteenth century concept of unitary psychosis [2], which mirrors the much wider application of RDoC to positing a similar transdiagnostic pathology for symptom domains across disorder categories. In RDoC, the hypothesized shared pathology within each domain is expected to differ only in severity across categories and even extend to subthreshold symptoms in healthy persons [1].

Conversely, if the biological processes producing symptoms differ across diagnostic categories, then research studies must again narrow their target subject groups based on diagnostic criteria. We recently reported such a circumstance for cognitive deficits within a psychosis group, in which we found that social cognition deficits were associated with distinct cellular abnormalities in schizophrenia compared to affective psychoses [3].

This study examined the cellular pathologies associated with symptoms across psychiatric categories by relating clinical symptoms to concentrations of whole-hippocampal neurometabolite biomarkers for cellular pathologies from proton MR spectroscopic imaging (^1^H MRSI) [4]. Associations of neurometabolite concentrations and symptoms were compared across individuals with non-psychotic affective disorders, psychotic disorders, and healthy controls, with the psychotic group also studied after stratification into schizophrenia and affective psychosis subgroups. The goal was to determine if specific symptom domains have similar cellular pathologies across diagnostic categories.

No previous MRS study of psychosis has compared associations of hippocampal metabolites and symptoms across different psychotic disorders, nor across psychotic and non-psychotic disorders. Instead, prior studies have almost always only compared group differences in mean metabolite levels between cases and controls, with highly variable results [5–12]. Limiting analyses to just mean metabolite levels may have reduced statistical power and lowered sensitivity to detect biological heterogeneity within psychosis [12]. Moreover, most studies have used single-voxel-spectroscopy, which yields limited area of coverage and spatial resolution, and usually examines just one side of hippocampus and misrepresents its irregular shape. We therefore used three-dimensional whole hippocampal multi-voxel MRS (3D ^1^H MRSI) to study the entire hippocampus with increased precision and spatial resolution.

## METHODS

### Human Subjects

This study prospectively recruited persons with psychosis (Psy) or non-psychotic affective disorders (NP-aff), irrespective of DSM-5 diagnosis, from treatment settings at Mount Sinai Hospital, and healthy controls (HC) from the community. Psy included cases with non-affective psychosis, i.e., schizophrenia (Scz), and affective psychosis (aff-P), the latter including participants with schizoaffective disorder and psychotic mood disorders. Cases were included if they had been on stable medication regimens for at least one month. HC had no personal or family history of psychosis, did not meet criteria for any DSM-5 diagnosis in the preceding two years, and were not taking psychiatric medications. Exclusion criteria included head trauma with sequalae, premorbid intellectual disability, suicide risk, seizures (other than medication-related) and MRI contraindications. All participants provided written informed consent for this Institutional Review Board approved study.

### Clinical Procedures

DSM-5 diagnoses were based on the Diagnostic Interview for Genetic Studies [13]. Symptoms were assessed with the Positive and Negative Syndrome Scale (PANSS) [14], from which we used the positive and negative symptom scales, and employed the PANSS pentagonal model [15] to examine factors for activation (behavioral dyscontrol), autistic preoccupation (thought disorder/disorganized behavior), and dysphoria (depression/anxiety) [16–18]. Depressive symptoms were more thoroughly assessed using the Hamilton Depression Scale (HAM-D) [19], and manic symptoms were measured with the Young Mania Rating Scale (YMRS) [20].

### MRI and Hippocampal Spectroscopic Imaging – Acquisition

Imaging was done in a 3 T whole-body MR scanner (Skyra, Siemens AG, Erlangen, Germany) using a transmit-receive head-coil (TEM3000, MR Instruments, Minneapolis, MN, USA). Subjects were placed head-first supine into the magnet, followed by sagittal *T_1_*-weighted magnetization-prepared rapid gradient-echo (MPRAGE) MRI: *T_E_*/*T_R_*/*T_I_* = 3.5/2150/1000 ms, 7° flip-angle, 160 slices 1 mm thick, 256×256 matrix, 256×256 mm^2^ field-of-view. These were reformatted into 192 axial, sagittal and coronal slices angled along the hippocampal long axis at 1 mm^3^ isotropic resolution, for ^1^H MRSI volume of interest (VOI) placement and tissue segmentation, as shown in Figure 1a,b [21].

**Figure 1:**
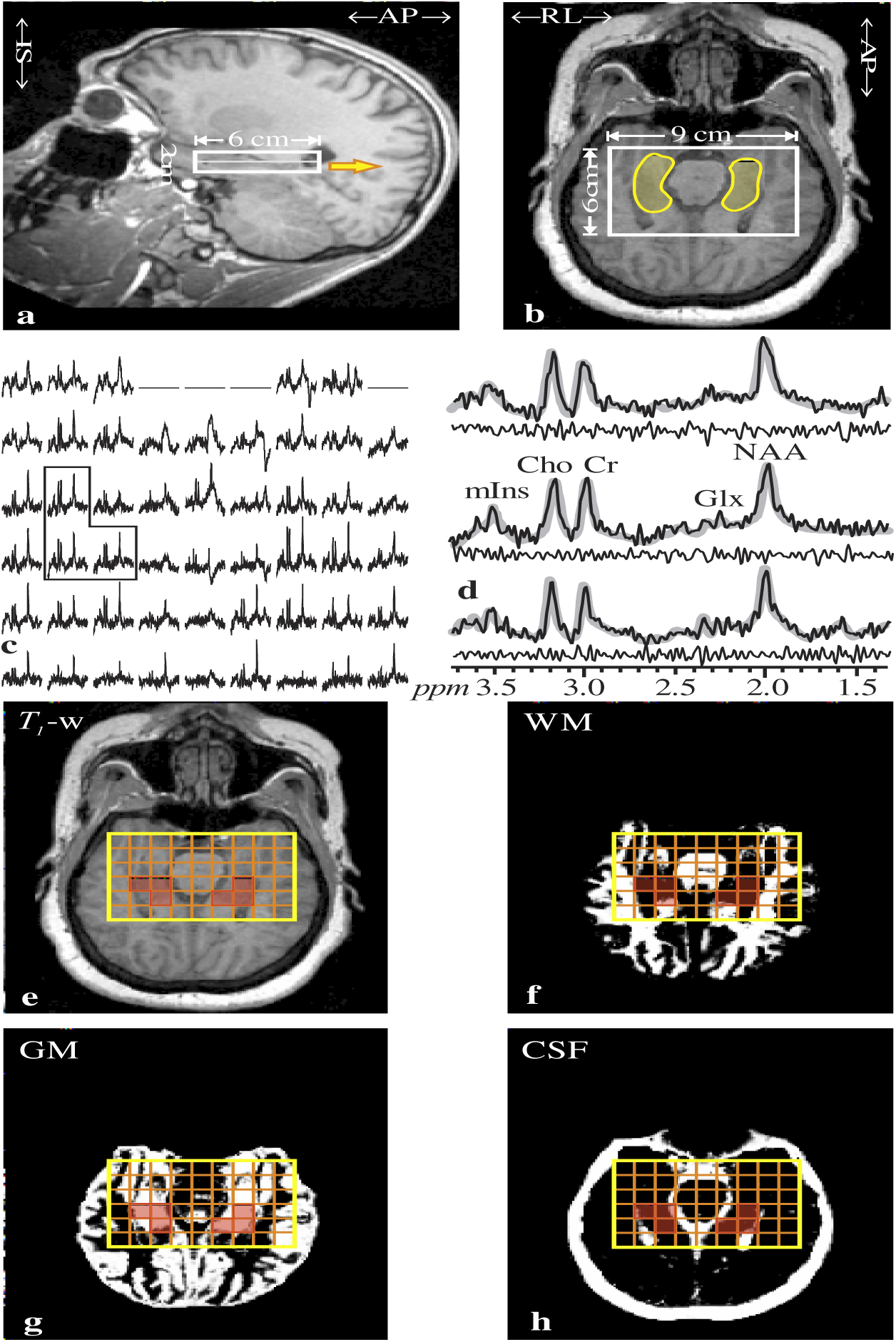
Imaging and 1H-MRSI position, size and analysis. **Note**. **Top:** Sagittal (**a**) and axial (**b**) *T*_1_-weighted MRI of a 22 year-old female with Scz, superimposed with the 9×6×2 cm^3 1^H MRSI VOI (thick white frames) and Frizoni guidelines based hippocampus outline (transparent yellow zones on **b**). Yellow arrow on a indicates the slice-level of **b**. **Center:** left - **c**: Real part of the 9×6 (LR×AP) axial ^1^H spectra matrix from the VOI slice on **b**. Spectra within the right hippocampus on a are marked by the dashed frame. Right – **d**: The spectra in that frame, expanded for detail (black lines) superimposed with the spectral-fit (gray, see Figure 2). Note the good signal-to-noise-ratio; spectral-resolution (8.1±3.0 Hz linewidth) from the (1 cm^3^) voxels; and fit fidelity, reflected by the “noise” (experimental – fit) residual below each spectrum. **Bottom: e – h**: Axial MRI (**e**), and its WM (**f**), GM (**g**), and CSF (**h**) probability maps, obtained with SPM12, superimposed with the ^1^H MRSI grid (orange matrix) and the bilateral hippocampus outline (transparent red). These are used by our software to identify hippocampal voxels and extract their GM spectroscopic content (corrected for CSF and WM partial volumes), as described in the Methods section.

A 6 cm anterior-posterior (AP) ×9 cm left-right (LR) ×2 cm inferior-superior (IS)=108 cm^3^ ^1^H MRSI VOI was then image guided over the bilateral hippocampus, as shown in Figure 1a,b, and excited with point resolved spectroscopy (PRESS: *TE*/*TR*= 120/1500 ms). This intermediate *TE* was a compromise to *(i)* attenuate short *T_2_* species; lipids and macromolecules’ signals from within the VOI, or those not sufficiently excluded by the selective PRESS pulses. This choice; *(ii)* suffer acceptable, 25-30% *T_2_* signal loss; and *(iii)* allows the *J*-coupled multiplets of Glx and mI to partially refocus, as shown in Figure 2. The VOI was encoded into two axial 1 cm thick slices, and 12×12 gradient-phase-encoded in their planes, as shown in Figure 1b [22, 23] to form (1.0 cm)^3^ voxels. The ^1^H MRSI signals were acquired for 256 ms at ±1 kHz bandwidth. At two averages the ^1^H MRSI was ~15 minutes and the entire protocol took under 40 minutes [21].

**Figure 2:**
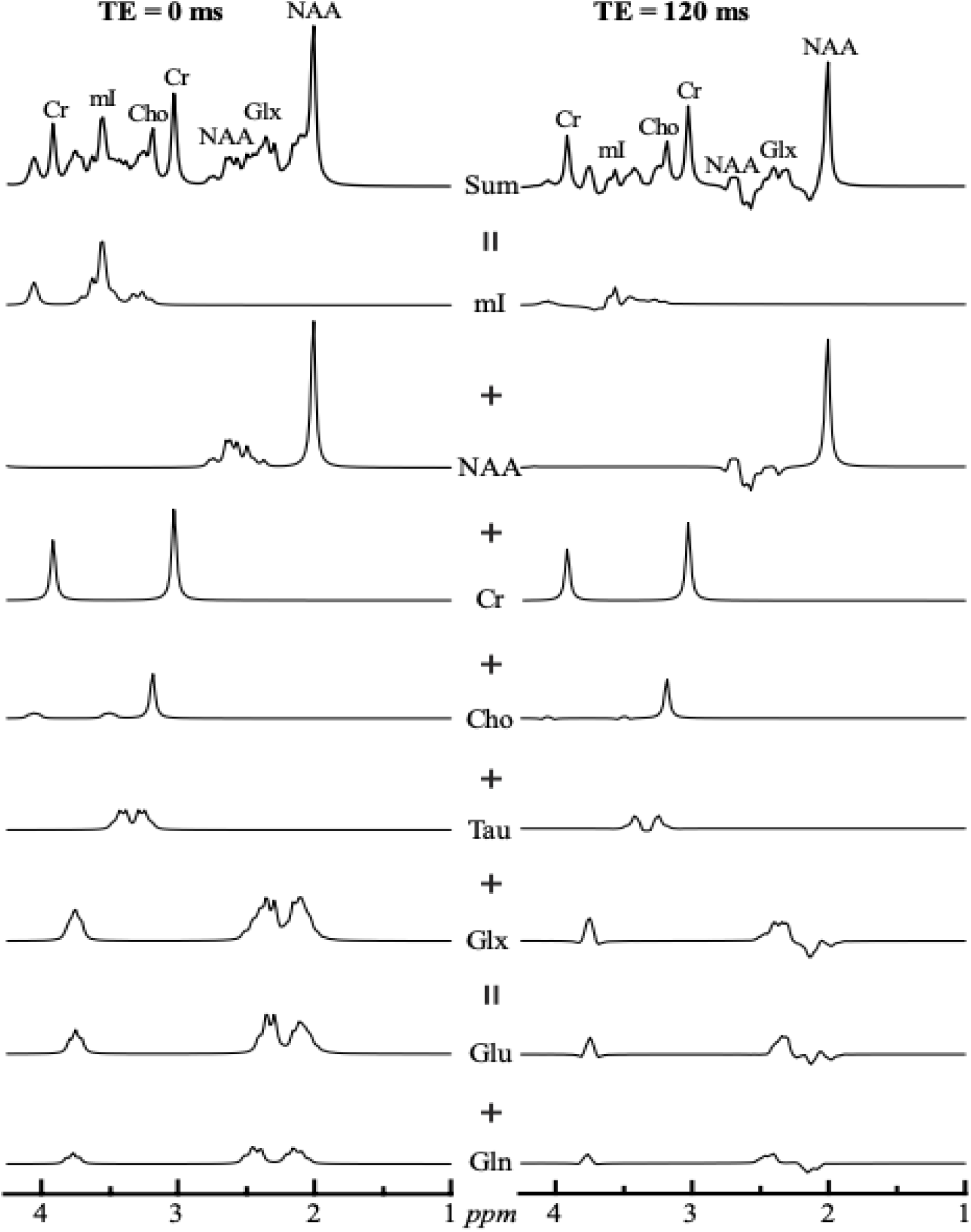
1H-MRSI spectral fitting model functions. Note. **Top**: Simulated ^1^H MRS spectra at *TE*=0 (left) and *TE*=120 ms (right), assuming metabolites’ *T*_2_=. These spectra were synthesized by summing the different simulated basis function of the individual metabolites’ (**below**), generated at *TE*=0 ms (left) and *TE*=120 ms (right), obtained by full density matrix-based spectral simulation for each metabolite in the GAVA-Simulation application using the sequence’s actual complex RF pulse waveforms and timings. The waveforms are weighted by their reported in vivo concentrations expected in the central nervous systems. NAA=10 mM, Cho=1 mM, Cr=6 mM, mI=6 mM, Glx=11.5 Mm (Glu=10 mM + Gln=1.5 mM), Taurine=4 mM. Note that (*i*) the J-coupled multiplets (Glx and mI) partially refocus, sufficiently to be distinct, even at *TE*=120 ms; (*ii*) the concordance of the full simulated *TE*=120 ms spectrum and the experimental spectra in Figure 1d.

### MRI and Hippocampal Spectroscopic Imaging – Metabolic Quantification

^1^H MRSI data were post-processed with in-house software (IDL 8.7.3, L3Harris Geospatial, Broomfield, CO). Residual water was removed from the signals in the time domain [24], the data magnetic-field drift corrected voxel-shifted to align the localization grid with the NAA VOI, zero-filled from 12×12 to 16×16 voxels in the slices’ planes and from 512 to 2048 in the time domain. The data was then Fourier transformed in the temporal, AP and LR directions and Hadamard reconstructed along the IS. Each spectrum was automatically corrected for frequency and zero-order phase shifts, as shown in Figure 1c,d.

The hippocampal metabolite concentrations used as biomarkers for cellular pathologies were *N*-acetyl-acetate (NAA) for neuronal integrity; Creatine (Cr) for energy metabolism; Choline (Cho) for myelin/membrane turnover; myo-inositol (mI) for glia, particularly astrocytes; and glutamate plus glutamine (Glx) for glutamatergic neurotransmission [4, 25]. Relative levels of each voxel’s NAA, Glx, Cr, Cho and mI were estimated from their spectral peak area using the SiTToolsFit spectral modeling package [26] (with the Glu+Gln (=Glx), Cho, Cr, *m*I, NAA and taurine model functions (Figure 2), as shown in Figure 1d. The relative levels were scaled into absolute concentrations against a 2 L reference sphere of known NAA, Cr, Cho and mI concentrations in water [27].

For tissue segmentation and ^1^H MRSI quantification, bilateral (left + right) hippocampal masks were traced by a trained neuroradiologist on the sagittal images, according to the Harmonized Protocol of the European Alzheimer’s Disease Consortium [28, 29] (see Figure 1b). The axial MRI were segmented into CSF, gray and white matter (GM, WM) masks with SPM12 [University College London, UK] [30], as shown in Figure 1f-h. In-house software (MATLAB 18, MathWorks, Framingham, MA) retained for analyses only voxels of: *(a)* at least 30% volume inside the hippocampus mask; *(b)<*30% CSF, *(c)* Cramer-Rao lower bounds <20% for any metabolite; and (*d*) 4 Hz<linewidths<13 Hz (both *(c)* and *(d)* are more stringent than the MRS consensus criteria described by Kreis and Wilson *et al.*) [31, 32], as shown in Figure 1e-h. The metabolites’ average GM concentrations in the retained (*N*>2) hippocampi voxels, was estimated by our software with linear regression, as described by Tal *et al.* [33].

### Statistical Analyses

ANOVA, Welch’s ANOVA, or Kruskal-Wallis test compared groups on age, score for each symptom domain, and each metabolite’s concentration. Chi-square test was used to compare the proportion of males to females across groups. For illness duration, Mann-Whitney U test compared Psy to NP-aff, and Kruskil-Wallis tests compared Scz, aff-P, and NP-aff. ANCOVA compared the groups on score for each symptom domain and each metabolite’s concentration while controlling for age and sex. Nonparametric Levene’s test compared the variance of each metabolite between groups. For each ANOVA/ANCOVA or equivalent non-parametric test and Chi-square test, *post-hoc* comparisons for each group combination were performed with correction for multiple comparisons to maintain α=.05. Within each group Spearman correlations assessed relationships between metabolite and symptom levels. For any significant correlations, Fisher’s r to z transformation was used to normalize the correlation outcome to allow for comparisons of correlation coefficients across groups. Post-hoc analysis compared significant correlations in Scz and aff-P with NP-aff and HC, and vice versa. Post-hoc analysis also examined intercorrelations between neurometabolites in schizophrenia to help with interpretation of findings in this group.

## RESULTS

The 81 participants included 33 with psychosis, including 20 with schizophrenia and 13 with affective psychosis, 22 participants with non-psychotic affective disorders, and 26 healthy controls. Some participants had missing values for mI and Glx due to decreased SNR for these metabolites, and one participant had missing values for the PANSS and HAM-D due to incompletion of these scales (see Supplementary Information).

Comparisons of demographics, symptom levels, and neurometabolite levels are compiled in Table 1. Age and sex were similar across groups, and disease duration did not differ across clinical groups.

**Table 1:** Demographics, Metabolite Concentrations, and Symptoms.

| Measure | Healthy Controls | Non-psychotic Affective Disorder | Psychosis | Schizophrenia | Affective Psychosis | Group Differences<sup>1</sup> |
| --- | --- | --- | --- | --- | --- | --- |
| **Demographics** |  |  |  |  |  |  |
| Age (Mean SD) | 32.15 (8.37) | 36.27 (11.82) | 35.79 (10.91) | 38.20 (2.40) | 32.08 (2.92) |  |
| Sex (males:females) | 8:18 | 6:16 | 17:16 | 12:8 | 5:8 |  |
| Duration of Illness (yr) |  | 15.55 (10.97) | 18.48 (12.61) | 20.05 (12.67) | 16.08 (12.64) |  |
| **PANSS Scores** |  |  |  |  |  |  |
| Positive Symptoms | 7.04 (.200) | 7.86 (1.39) | 15.88 (6.19) | 18.35 (1.41) | 12.08 (1.01) | Psy, Scz aff-P > HC, NP-aff Scz > aff-P |
| Negative Symptoms | 7.00 (0.00) | 9.41 (3.96) | 11.30 (4.85) | 12.45 (1.29) | 9.54 (.606) | Psy, Scz > HC |
| Activation | 6.04 (.200) | 6.59 (1.22) | 8.36 (2.98) | 8.35 (.719) | 8.38 (.747) | Psy, Scz, aff-P > HC, NP-aff |
| Autistic | 6.00 (.000) | 6.59 (1.40) | 9.85 (3.06) | 11.20 (.618) | 7.77 (.632) | Psy, Scz > HC, NP-aff Scz > aff-P > HC |
| Preoccupation | 5.12 (.440) | 11.05 (3.74) | 8.94 (4.25) | 7.95 (.884) | 10.46 (1.22) | Psy, Scz, aff-P, NP-aff > HC Scz > NP-aff |
| Dysphoria |  |  |  |  |  |  |
| YMRS Score | .270 (.604) | .409 (1.01) | 3.00 (7.05) | 4.20 (1.98) | 1.15 (.421) | Scz > HC |
| HDRS Score | .720 (1.28) | 8.55 (6.06) | 4.88 (5.61) | 4.30 (1.37) | 5.77 (1.34) | NP-aff > Psy > HC Aff-P > HC |
| **Metabolite Concentration** |  |  |  |  |  |  |
| *Mean (SD) millimolar* |  |  |  |  |  |  |
| NAA | 6.65 (1.49) | 6.13 (1.68) | 6.64 (1.36) | 6.69 (.332) | 6.57 (.331) |  |
| Cr | 7.47 (1.61) | 7.44 (1.10) | 7.05 (2.17) | 7.56 (.474) | 6.26 (.581) |  |
| Cho | 2.48 (.630) | 2.48 (.476) | 2.51 (.696) | 2.44 (.161) | 2.62 (.186) |  |
| mI | 4.84 (.997) | 4.74 (1.36) | 4.21 (1.52) | 4.02 (.386) | 4.60 (.643) |  |
| Glx | 7.82 (2.06) | 6.86 (2.31) | 6.73 (2.97) | 7.54 (1.01) | 5.39 (.888) |  |

Note. <sup>1</sup>Only significant differences when controlling for age and sex noted in this column; HC: Healthy controls, Psy: Psychosis, NP-aff: Non-psychotic affective disorder; Scz: schizophrenia; aff-P: Affective Psychosis.

### 1. Comparisons of Symptom Scores Across Groups

Group comparisons (Table 1) of symptoms, both before and after stratification of Psy into psychosis subgroups, showed significant differences between groups for positive and negative symptoms, activation, autistic preoccupation, dysphoria, and depression, with large effect sizes (ranging from η2=.175 to .503), unaltered when adjusting for age and sex (p’s<.001). Post-hoc comparisons showed that all of these symptoms were increased in Psy compared to HC (*p*’s≤.002). These symptoms were also increased in the psychosis subgroups (Scz and aff-P) compared to HC before adjustment for age and sex, except for depression in Scz (*p’s*<.008). The results were the same after controlling for age and sex, except for an increase in significance for manic symptoms in Scz (p=.007) and loss of significance for negative symptoms in aff-P. The latter may reflect the combined effects of age and sex on negative symptoms in affective psychosis, as post-hoc analysis showed a similar result when only controlling for age and for sex-stratified comparisons. When comparing the two psychosis subgroups, Scz had higher positive symptoms and autistic preoccupation than aff-P (*p’s*<.008).

Compared to NP-aff, Psy showed higher positive symptoms, activation, and autistic preoccupation (*p’s* ≤ 01), and decreased depression (p=.016), all unaltered by adjusting for age and sex. Similarly, compared to NP-aff, Scz and aff-P showed increased positive symptoms and activation, and Scz also showed elevated autistic preoccupation and reduced dysphoria (*p’s*<.008). Compared to HC, NP-aff showed increased depression and dysphoria (*p’s*<.01).

### 2. Comparisons of Neurometabolite Concentrations Across Groups

All groups were similar in mean levels of each of the hippocampal neurometabolites, both before and after controlling for age and sex (Table 1). Groups were also similar in the variance of each neurometabolite measure.

### 3. Associations of Neurometabolite Concentrations and Symptom Scores

Despite similar neurometabolite levels between groups, we found significant within-group associations of neurometabolite levels with symptom scores (Table 2 and Supplementary Tables S1-S12). Significant findings involving very weak correlations (<.2) are omitted in the text but shown in the tables.

a. **HC:** In HC, but not the clinical groups, higher levels of the glia biomarker (mI) were moderately associated with increased manic (ρ=.49, p=.025) and depressive symptoms (ρ=.55, p=.012), with the latter association opposite in Psy (p=.007) and Scz (p=.032).
b. **NP-aff:** In NP-aff, lower concentrations of the biomarker for cellular energy (Cr) were moderately associated with increased activation (ρ=−.46, p=.030) and dysphoria scores (ρ=−.48, p=.025). Also, reduced levels of the glia biomarker (mI) were moderately related to increased negative symptoms (ρ=−.67, p=.005) and autistic preoccupation (p=−.55, p=.029).
c. **Psy Group (combined Scz and aff-P subgroups):** In Psy, activation was moderately associated with mI (ρ=−.69, p<.001), and moderately related to Cr (ρ=−.43, p=.012) and the markers for neural integrity (NAA) (ρ=−.42, p=.015) and neural excitation (Glx) (ρ=−.51, p=.044), with the association with mI opposite in HC (p<.001). Also in Psy, lower Cr was furthermore moderately associated with increased depression (ρ=−.42, p=.016) and dysphoria (ρ=−.45, p=.008).
d. **Scz:** The associations of increased activation with reduced metabolite levels in Psy were mostly explained by Scz, in which activation was moderately to strongly related to lower levels of all five metabolites; mI (ρ=−.84, p<.001), NAA (ρ=−.54, p=.014), Cr (ρ=−.50, p=.024), Glx (ρ=−.66, p=.037), and Cho (the membrane turnover marker) (ρ=−.45, p=.047). The association with Cho in Scz was opposite in aff-P and HC (p’s<.05), and the association with mI opposite in HC (p<.001). Decreased Cho in Scz was additionally moderately related to increased positive symptoms (ρ=−.52, p=.018), autistic preoccupation (ρ=−.45, p=.046), and dysphoria (ρ=−.45, p=.048), with the associations with positive symptoms and autistic preoccupation opposite in aff-P (p’s<.05). Elevated autistic preoccupation in Scz was also moderately related to lower mI (ρ=−.55, p=.044). Cr in Scz was additionally moderately associated with dysphoria (ρ=−.61, p=.005) and depressive symptoms (ρ=−.45, ρ=.0496), which explained the associations of the same measures in Psy. Also in Scz, increased depression was additionally moderately related to reduced NAA (ρ=−.45, p=.047), which was opposite in NP-aff (p=.016).
e. **aff-P:** Similarly to NP-aff, in aff-P decreased mI was strongly associated with increased negative symptoms (ρ=−.77, p=.042) and autistic preoccupation (ρ=−.78, p=.040). Also in aff-P, higher Cho was moderately related to increased manic symptoms (ρ=.65, p=.015), which was opposite in HC and NP-aff (p’s<.05).
f. **Comparing symptom domains across groups based on neurometabolite profiles:** For activation, a process impacting all cellular components was implicated in schizophrenia, but only reduced energy metabolism was suggested in non-psychotic affective cases. For mood symptoms, increased glia was suggested for healthy controls, whereas increased membrane turnover was related to mania in affective psychosis, and reduced neural integrity and energetics were related to depression in schizophrenia. Affective groups with or without psychosis both showed less glia in association with negative symptoms and autistic preoccupation, the latter also in schizophrenia, although in schizophrenia autistic preoccupation was also associated with reduced membrane turnover. Dysphoria was associated with reduced energy metabolism in both non-psychotic affective cases and schizophrenia, but also with reduced membrane turnover in schizophrenia. Positive symptoms were only related to cellular abnormalities in schizophrenia, specifically reduced membrane turnover.

**Table 2:**
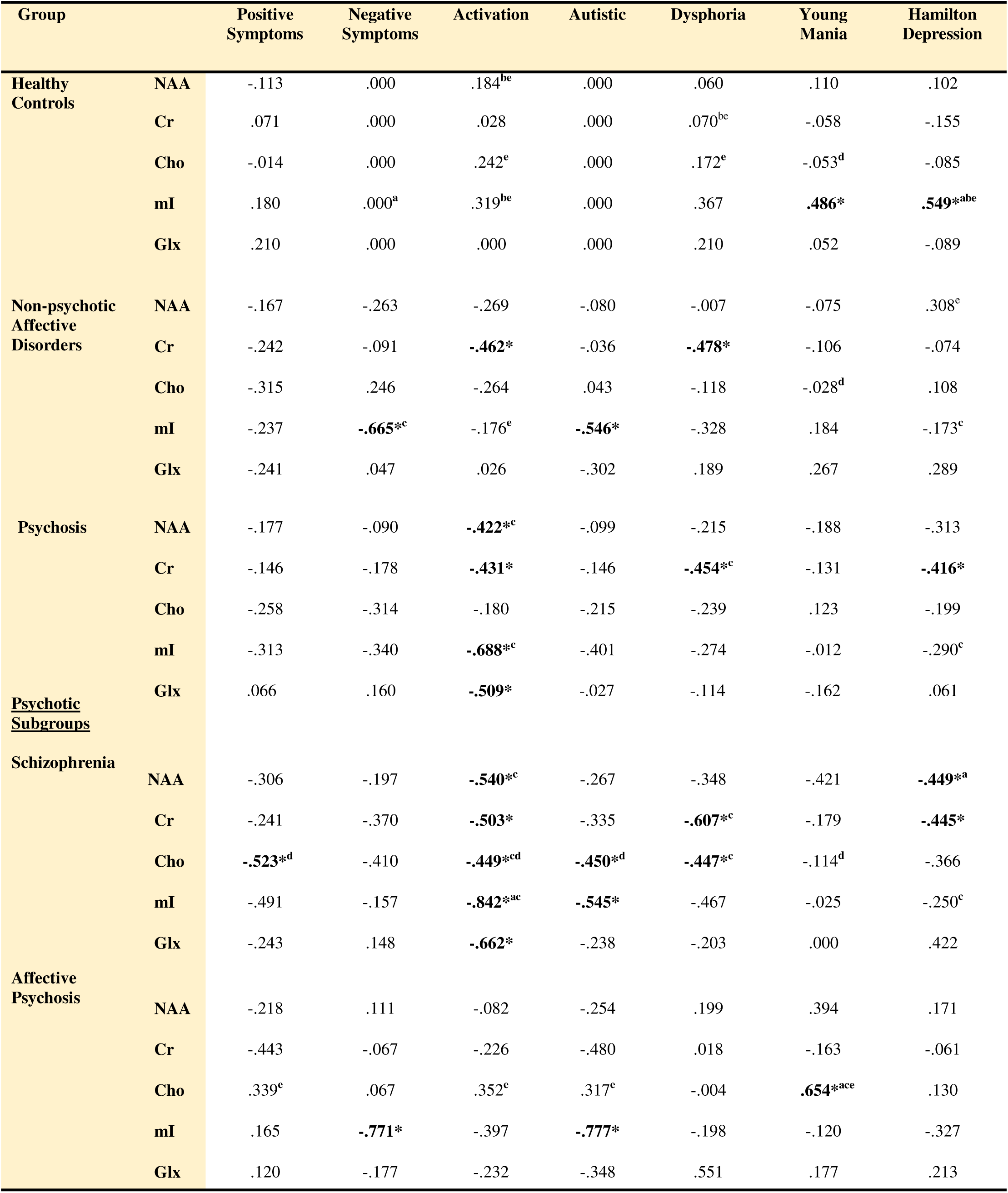
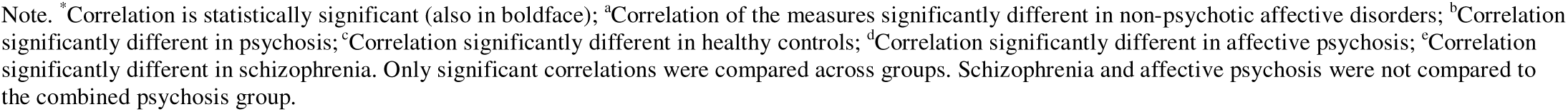
Correlations of Each Metabolite Concentration and Symptom Score by Diagnostic Category.

| Group |  | Positive Symptoms | Negative Symptoms | Activation | Autistic | Dysphoria | Young Mania | Hamilton Depression |
| --- | --- | --- | --- | --- | --- | --- | --- | --- |
| Healthy Controls | NAA | -.113 | .000 | .184<sup>be</sup> | .000 | .060 | .110 | .102 |
|  | Cr | .071 | .000 | .028 | .000 | .070<sup>be</sup> | -.058 | -.155 |
|  | Cho | -.014 | .000 | .242<sup>e</sup> | .000 | .172<sup>e</sup> | -.053<sup>d</sup> | -.085 |
|  | mI | .180 | .000<sup>a</sup> | .319<sup>be</sup> | .000 | .367 | **.486\*** | **.549\*<sup>abe</sup>** |
|  | Glx | .210 | .000 | .000 | .000 | .210 | .052 | -.089 |
| Non-psychotic Affective Disorders | NAA | -.167 | -.263 | -.269 | -.080 | -.007 | -.075 | .308<sup>e</sup> |
|  | Cr | -.242 | -.091 | **-.462\*** | -.036 | **-.478\*** | -.106 | -.074 |
|  | Cho | -.315 | .246 | -.264 | .043 | -.118 | -.028<sup>d</sup> | .108 |
|  | mI | -.237 | **-.665\*<sup>c</sup>** | -.176<sup>e</sup> | **-.546\*** | -.328 | .184 | -.173<sup>c</sup> |
|  | Glx | -.241 | .047 | .026 | -.302 | .189 | .267 | .289 |
| Psychosis | NAA | -.177 | -.090 | **-.422\*<sup>c</sup>** | -.099 | -.215 | -.188 | -.313 |
|  | Cr | -.146 | -.178 | **-.431\*** | -.146 | **-.454\*<sup>c</sup>** | -.131 | **-.416\*** |
|  | Cho | -.258 | -.314 | -.180 | -.215 | -.239 | .123 | -.199 |
|  | mI | -.313 | -.340 | **-.688\*<sup>c</sup>** | -.401 | -.274 | -.012 | -.290<sup>c</sup> |
|  | Glx | .066 | .160 | **-.509\*** | -.027 | -.114 | -.162 | .061 |
| <u>Psychotic Subgroups</u> |  |  |  |  |  |  |  |  |
| Schizophrenia | NAA | -.306 | -.197 | **-.540\*<sup>c</sup>** | -.267 | -.348 | -.421 | **-.449\*<sup>a</sup>** |
|  | Cr | -.241 | -.370 | **-.503\*** | -.335 | **-.607\*<sup>c</sup>** | -.179 | **-.445\*** |
|  | Cho | **-.523\*<sup>d</sup>** | -.410 | **-.449\*<sup>cd</sup>** | **-.450\*<sup>d</sup>** | **-.447\*<sup>c</sup>** | -.114<sup>d</sup> | -.366 |
|  | mI | -.491 | -.157 | **-.842\*<sup>ac</sup>** | **-.545\*** | -.467 | -.025 | -.250<sup>c</sup> |
|  | Glx | -.243 | .148 | **-.662\*** | -.238 | -.203 | .000 | .422 |
| Affective Psychosis | NAA | -.218 | .111 | -.082 | -.254 | .199 | .394 | .171 |
|  | Cr | -.443 | -.067 | -.226 | -.480 | .018 | -.163 | -.061 |
|  | Cho | .339<sup>e</sup> | .067 | .352<sup>e</sup> | .317<sup>e</sup> | -.004 | **.654\*<sup>ace</sup>** | .130 |
|  | mI | .165 | **-.771\*** | -.397 | **-.777\*** | -.198 | -.120 | -.327 |
|  | Glx | .120 | -.177 | -.232 | -.348 | .551 | .177 | .213 |

### 4. Post-hoc Intercorrelations of neurometabolites in Scz

There were moderate to very strong positive correlations between all neurometabolites (NAA, Cr, Cho, mI, and Glx), all significant except for trend level associations of NAA with Cr, as well as Cho with Glx (Figure 3 and Supplementary Table S13).

**Figure 3:**
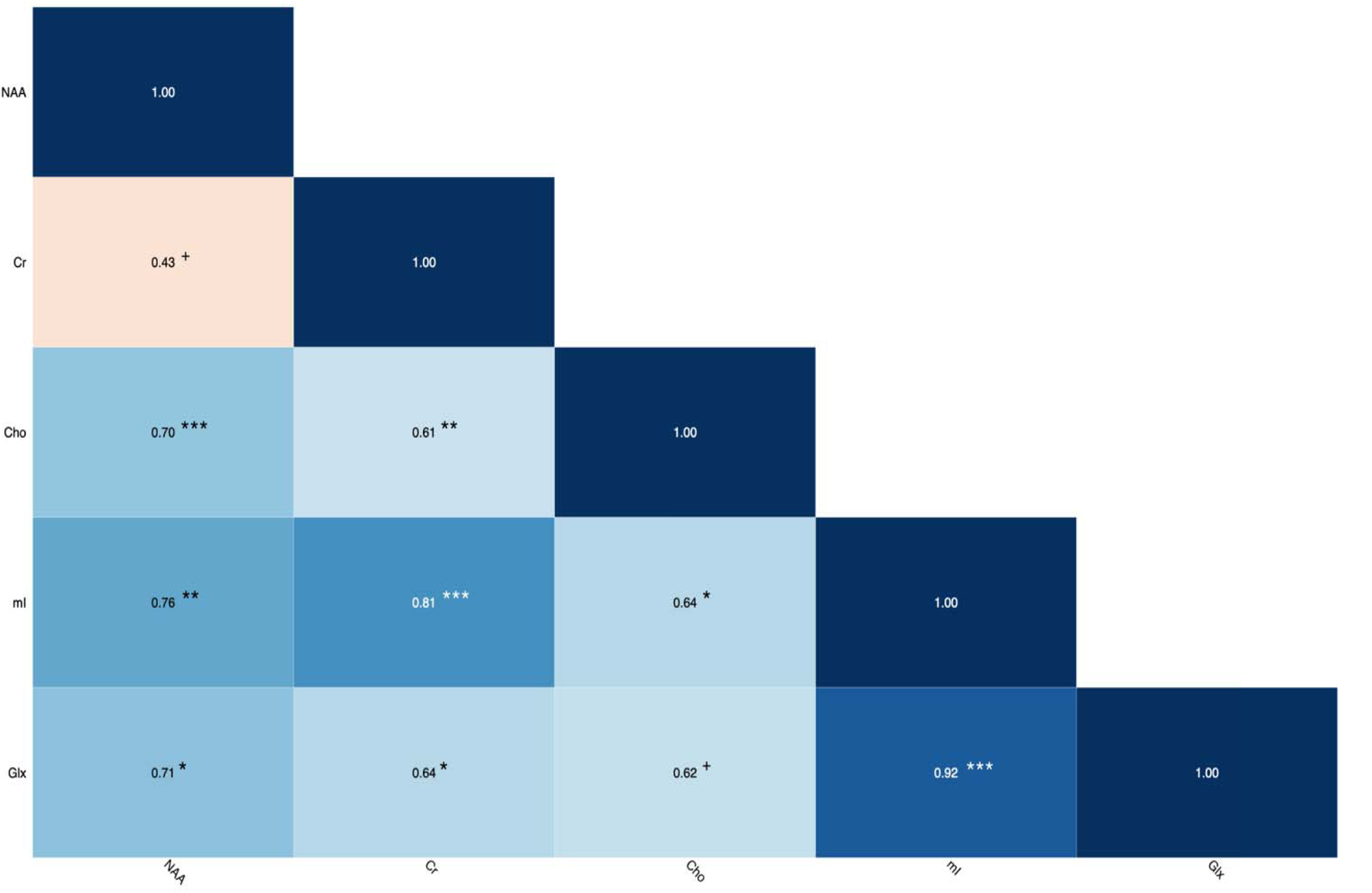
Intercorrelations Between Neurometabolites in Schizophrenia. Note. The correlation matrix heatmap shows the Spearman correlation coefficient values (ρ) for all combinations of neurometabolites (NAA, Cr, Cho, mI, Glx), ranging from 0.0 to 1.0, with 1.0 indicating a perfect positive correlation between variables and 0.0 indicating no correlation between variables. Statistical significance is indicated by asterisks (*** for p<.001; ** for p<.01; * for p <.05), and trend towards significance indicated by plus symbol (+). Note: Moderate to very strong positive correlations were shown between all neurometabolites, all significant (p’s<.05), except for trend level correlations of NAA with Cr (p=−.058), and of Cho with Glx (p=.055).

## DISCUSSION

This analysis suggests distinct cellular underpinnings across psychotic, non-psychotic affective, and healthy participants for the PANSS activation factor and mood symptoms (YMRS and HAM-D). On the other hand, mostly similar cellular abnormalities were shown for the autistic preoccupation factor and negative symptoms, and both differences and similarities were shown for the dysphoria factor. Positive symptoms were associated with cellular abnormalities only in the schizophrenia group. Symptom severity levels were largely in line with the literature, as psychosis had higher levels of all PANSS symptom scales and factors than healthy controls, as well as higher positive symptoms, autistic preoccupation, and activation, and lower depression than non-psychotic affective cases.

The activation factor was related to reductions across all metabolite biomarkers for cellular pathologies in schizophrenia, but only to reduced Cr (energy use) in non-psychotic affective cases. The widespread deficits in schizophrenia suggest a process jointly impacting all cellular components, including neural integrity, energetics, membrane turnover, glia, and glutaminergic transmission, which is supported by post-hoc results of moderate to very strong intercorrelations between all metabolites in this group. A systemic condition may explain this abnormal metabolite profile, based on our prior linkage of the activation factor to systemic inflammatory conditions and medical comorbidity [34].

Another plausible pathway is one centered around neurodevelopmental glial deficits, as reflected by the reductions in Cho (membrane turnover) and mI (glia), which may indicate hypomyelination and astrocyte deficits, respectively. The former aligns with deficits in white matter, oligodendrocytes, and myelin in schizophrenia studies, as well as reduced expression of genes/proteins related to oligodendrocytes and myelin [35–37]. The white matter deficits are linked to functional dysconnectivity in the illness [38], consistent with less myelin impairing functional connectivity by degrading signal transduction [39]. Like oligodendrocytes, astrocytes serve critical neuroprotective functions, and astrocytes also play a crucial role in directing synaptogenesis [40]. Less synaptogenesis during neurodevelopment alongside the excess synaptic pruning associated with schizophrenia [41] may contribute to the reduced neuropil implicated in the illness (e.g., decreased synaptic [42] and dendritic spine density [43]).

The reduced neural integrity (NAA) and glutaminergic transmission (Glx) may reflect neurodegeneration that is caused or exacerbated by the loss of glia and their neuroprotective effects, with the lower Glx specifically reflecting reduced neuropil density of the glutaminergic system [44]. As such, less Glx may also reflect the inability of neurons to synthesize glutamate without the glutamine produced by astrocytes and/or a reduction in astrocytic coordination of neuronal excitatory/inhibitory balance [45]. In this context, the astrocyte loss could be of relevance to long-standing theories of schizophrenia pathogenesis which posit glutaminergic dysregulation due to NMDA receptor hypofunction [46]. Such theories usually predict excess glutamate release, and so our reduced Glx finding could reflect a compensatory response to excitotoxicity that attempts to hinder neuronal damage, with astrocyte dysfunction further exacerbating this cycle via reduced glutamate uptake. Reduced Glx may therefore indicate an overall glutamate loss rather than synaptic glutamate loss specifically [47], which is consistent with the Glx signal owing primarily to intracellular rather than extracellular glutamate [48, 49]. As for the reduced Cr, lower energy use is plausible given the widespread metabolic deficits we found, although less Cr could more simply reflect the glial loss in that Cr has a twofold to threefold greater concentration in glia than neurons [50].

Reduced Cho was involved in a broad array of symptoms in schizophrenia, not only activation, but also positive symptoms, autistic preoccupation, and dysphoria, suggesting a possible role for hypomyelination across multiple symptom domains. The white matter deficits in the illness are frequently linked to positive symptoms [51]. Moreover, white matter changes [52, 53] and functional connectivity deficits [54, 55] are also related to items of the activation factor in schizophrenia, like hostility/aggression and poor impulse control. Hypomyelination may thus link positive symptoms, activation, and other symptoms to neurodevelopmental etiologies in schizophrenia (e.g., genetic vulnerability and/or maternal/fetal insults to the brain) [35, 37], which could more broadly account for the glial deficits proposed for activation (as above). For example, microglial activation and inflammation in the brain during early development [56–58] could cause reduced glial progenitor cell proliferation and differentiation, and in effect delayed maturation and reduced numbers of oligodendrocytes and astrocytes [56].

In our non-psychotic affective cases, less energy use (Cr) was linked to both the activation and dysphoria factors, the latter which includes items for some features associated with depression. Accordingly, lower energy use is implicated in the pathogenesis of mood disorders [59]. Reduced cellular energy in depression is related to mitochondrial dysregulation and reduced glucose availability [60, 61], as well as lower levels of phosphocreatine/creatine and lactate [60], which are critical for rapid ATP regeneration in neurons and glia. These abnormalities in depression may be caused or exacerbated by chronic stress and glucocorticoid excess [60, 61], a pathway linked to glutamatergic dysregulation, neuron loss, and impairments with neuroplasticity [61–63].

The differences in associations of metabolites with activation between schizophrenia and non-psychotic affective cases are noteworthy. Activation is a transdiagnostic domain seen in both psychotic and affective disorders [64]. It includes items on hostility, poor impulse control, excitement, and uncooperativeness, and has been implicated in aggressive behaviors, indicating a need for management. Activation captures behavioral dyscontrol and increased psychomotor activity and agitation, distinct from positive, negative, dysphoria (depression/anxiety), and autistic preoccupation (thought disorder/disorganized behavior) factors [16–18].

Cellular abnormalities for HAM-D and YMRS mood symptoms also differed across groups, particularly between the healthy and psychotic groups. The link of subclinical mood symptoms with higher hippocampal mI in otherwise healthy subjects may accord with results suggesting gliosis as a vulnerability marker for mood disorders in certain cases [65, 66]. In contrast, depressive symptoms in schizophrenia were linked to markers for reduced neural integrity and energy use, which may be of relevance to the reduced neuropil associated with the illness [41]. It could be that less neuropil leads to less energy demand, or conversely that less energy production can’t support sufficient synaptic connections and arborization [41]. Manic symptoms in affective psychosis were related to increased membrane turnover (Cho), possibly reflecting demyelination, which is consistent with reduced white matter integrity in affective psychoses [67–70]. The most common cause of demyelination is inflammation, and accordingly studies show increased levels of proinflammatory cytokines [71–73], autoimmune disease [74], and infection [75] in psychoses. In the brain, hippocampus is specifically linked to inflammation in psychoses by gene-expression studies [76].

Negative symptoms and autistic preoccupation were linked to similar cellular abnormalities across affective disorders with and without psychosis, specifically reduced mI. In line with this finding, reduced hippocampal glia is consistently reported in mood disorders and may contribute to mood symptoms in these illnesses via disruption of neuronal support and synaptic function [77–79]. The same cellular pathologies may also contribute to negative symptoms and autistic preoccupation in mood disorders both with and without psychosis.

As in the affective groups, autistic preoccupation was also related to reduced mI in schizophrenia, although in the latter it was additionally related to reduced Cho, which may align with the glial deficits proposed for activation in this group (as above). There could be separable pathologies for autistic preoccupation in schizophrenia, one shared with both affective groups and another distinct. Alternatively, entirely distinct pathologies may underpin autistic preoccupation between the schizophrenia and affective groups. These possibilities are further supported by our finding that the correlation of autistic preoccupation and Cho in schizophrenia was opposite in affective psychosis. These competing explanations may also apply to our findings of both overlap and differences in underpinnings for dysphoria, as it was related to reduced energetics in both non-psychotic affective cases (as above) and schizophrenia, but also to less membrane turnover in the latter.

Importantly, stratifying the psychosis group into subgroups showed that the associations observed in the combined psychosis group were mostly explained by schizophrenia, with distinct associations in affective psychosis. Moreover, before stratification the substantial involvement of reduced Cho (membrane turnover) was entirely occluded, as was the fact that the associations of Cho with symptoms in schizophrenia were consistently opposite in affective psychosis.

We found differential associations of symptoms and metabolites between groups despite no differences in mean metabolite levels. Thus, it may be that metabolite levels represent pathology by clinical phenomenon (e.g., specific symptoms or cognitive deficits) rather than simply by total metabolite levels. Also, certain subsets of cases that in part cluster together based on form of psychosis may explain the associations, supported by schizophrenia but not affective psychosis mostly driving the associations in the combined psychosis group.

The transdiagnostic RDoC model is not supported by our findings of distinct underpinnings for activation and mood symptoms across groups, although it is consistent with the overlap we found for autistic preoccupation and negative symptoms. Therefore, our findings suggest a far more nuanced picture than RDoC predicts.

This is the first MRS study of hippocampus in psychosis to: (1) test relationships of symptoms and metabolites across psychotic and non-psychotic groups; (2) test relationships across different psychoses; (3) examine PANSS factors outside of the standard positive and negative symptom domains. Additionally, the 3D whole hippocampus multi-voxel MRS we used was a significant improvement on the single-voxel techniques used in most past studies, yielding complete coverage of the irregularly-shaped hippocampus with increased spatial resolution.

Despite these strengths, our study had limitations. First, although the overall sample was sufficiently large, more modest numbers were observed for some analyses in the affective psychosis and schizophrenia subgroups. Additionally, while focusing on hippocampus was well justified, other brain areas which may reveal different associations were not included in the spectroscopic VOI in order to shorten acquisition time and minimize contamination by extraneous signals.

In conclusion, this novel MRSI study of neurometabolite markers for symptoms found both differences and similarities across psychotic, non-psychotic affective, and healthy groups, as well as across affective and non-affective psychoses. The differences taken alongside schizophrenia but not affective psychosis mostly driving the cellular pathology for symptoms in the full psychosis group support both categorizing disorders and stratifying different psychoses in research as opposed to fully abandoning diagnostic labels in favor of transdiagnostic approaches. As such, our results suggest that the common research practice of intermixing all psychoses may contribute to the lack of effective treatments.

## ACKNOWLEDGEMENTS

This work was supported by NIMH 1R01MH110418-01A1 (DM), NIBIB P41 EB017183 (OG), and NIBIB U24 EB02898 (OG).

