## Supplementary Information for "Do Symptom Domains Have Similar Cellular Underpinnings Across Psychiatric Diagnoses: Evidence from 3D Hippocampal MR Spectroscopy"

***Supplementary Materials and Methods***

There was some missing data for certain participants. There were some missing values across all of the groups for MRS measurements of mI and Glx because the peaks for mI and Glx in the MRS spectrum are much smaller than the peaks for the other metabolites measured, and thus SNRs for mI and Glx are sometimes lower than acceptable. Additionally, there was one participant from the healthy control group who didn’t complete the PANSS or the Hamilton Depression Rating Scale and therefore didn’t have values for PANSS positive symptoms, negative symptoms, activation, autistic preoccupation, dysphoria, and Hamilton Depression Rating Scale. Compared to the full sample of 33 persons with psychosis (Psy), including 20 with schizophrenia (Scz) and 13 with affective psychosis (aff-P), 26 healthy controls (HC), and 22 persons with non-psychotic affective disorders (NP-aff): in Psy there were 21 participants with mI values, including 14 with Scz and 7 with aff-P, and 16 participants with Glx values, including 10 with Scz and 6 with aff-P; in NP-aff there were 16 participants with mI values and 16 with Glx values; in HC there were 21 participants with mI values and 21 with Glx values; also in HC there were 25 participants with values for clinical measures other than The Young Mania Rating Scale (YMRS); for correlation analyses in HC of mI with any of the clinical measures other than YMRS there were 20 participants, and for correlations of mI with YMRS in HC there were 21 participants; for correlation analyses in HC of Glx with any of the clinical measures other than YMRS there were 20 participants and for correlations of Glx and YMRS in HC there were 21 participants.

***Supplementary Results***

**Table S1: Significant correlations (p** ≤ **.05) in Psychosis of Metabolites with Symptoms**

| **Variables correlated** | **Sample size** | **Correlation coefficient (rho)** | **P value (two-tailed)** |
| --- | --- | --- | --- |
| Activation - NAA | 33 | rho = -.422 | .015 |
| Activation - Cr | 33 | rho = -.431 | .012 |
| Dysphoria - Cr | 33 | rho = -.454 | .008 |
| Activation - mI | 21 | rho = -.688 | <.001 |
| Activation - Glx | 16 | rho = -.509 | .044 |
| Depressive Symptoms - Cr | 33 | rho = -.416 | .016 |

**Table S2: Significant correlations (p** ≤ **.05) in Healthy Controls of Metabolites with Symptoms**

| **Variables correlated** | **Sample size** | **Correlation coefficient (rho)** | **P value (two-tailed)** |
| --- | --- | --- | --- |
| Depressive Symptoms - mI | 20 | rho = .549 | .012 |
| Manic Symptoms - mI | 21 | rho = .486 | .025 |

**Table S3: Significant correlations (p** ≤ **.05) in Non-psychotic Affective Disorder of Metabolites with Symptoms**

| **Variables correlated** | **Sample size** | **Correlation coefficient (rho)** | **P value (two-tailed)** |
| --- | --- | --- | --- |
| Negative Symptoms - mI | 16 | rho = -.665 | .005 |
| Activation - Cr | 22 | rho = -.462 | .030 |
| Dysphoria - Cr | 22 | rho = -.478 | .025 |
| Autistic Preoccupation - mI | 16 | rho = -.546 | .029 |

**Table S4: Significant differences (p** ≤ **.05) in correlation coefficients: Healthy Controls (HC) vs. Psychosis (Psy)**

| **Variables correlated** | **HC correlation coefficient (rho)** | **HC sample size** | **Psy correlation coefficient (rho)** | **Psy sample size** | **Z statistic** | **P value (two-tailed)** |
| --- | --- | --- | --- | --- | --- | --- |
| Activation - NAA | rho = .184 | 25 | rho = -.422 | 33 | 2.27 | .023 |
| Activation - mI | rho = .319 | 20 | rho = -.688 | 21 | 3.47 | <.001 |
| Depressive Symptoms - mI | rho = .549 | 20 | rho = -.290 | 21 | 2.71 | .007 |
| Dysphoria - Cr | rho = .070 | 25 | rho = .478 | 33 | 1.99 | .047 |

**Table S5: Significant differences (p** ≤ **.05) in correlation coefficients: Healthy Controls (HC) vs. Non-psychotic affective Disorder (**NP-aff**)**

| **Variables correlated** | **HC correlation coefficient (rho)** | **HC correlation coefficient (rho)** | **NP-aff correlation coefficient (rho)** | **NP-aff sample size** | **Z statistic** | **P value (two-tailed)** |
| --- | --- | --- | --- | --- | --- | --- |
| Negative Symptoms - mI | rho = .000 | 19 | rho = -.665 | 16 | 2.15 | .032 |
| Depressive Symptoms - mI | rho = .543 | 20 | rho = -.173 | 16 | 2.15 | .032 |

**Table S6: Significant correlations (p** ≤ **.05) in Schizophrenia of Metabolites with Symptoms**

| **Variables correlated** | **Sample size** | **Correlation coefficient (rho)** | **P value (two-tailed)** |
| --- | --- | --- | --- |
| Positive Symptoms - Cho | 20 | rho = -.523 | .018 |
| Activation - NAA | 20 | rho = -.540 | .014 |
| Activation - Cr | 20 | rho = -.503 | .024 |
| Activation - Cho | 20 | rho = -.449 | .047 |
| Autistic Preoccupation - Cho | 20 | rho = -.450 | .046 |
| Dysphoria - Cr | 20 | rho = -.607 | .005 |
| Dysphoria - Cho | 20 | rho = -.447 | .048 |
| Activation - mI | 14 | rho = -.842 | <.001 |
| Autistic Preoccupation - mI | 14 | rho = -.545 | .044 |
| Activation - Glx | 10 | rho = -.662 | .037 |
| Depressive Symptoms - NAA | 20 | rho = -.449 | .047 |
| Depressive Symptoms - Cr | 20 | rho = -.445 | .0496 |

**Table S7: Significant correlations (p** ≤ **.05) in Affective Psychosis of Metabolites with Symptoms**

| **Variables correlated** | **Sample size** | **Correlation coefficient (rho)** | **P value (two-tailed)** |
| --- | --- | --- | --- |
| Autistic Preoccupation - mI | 7 | rho = -.777 | .040 |
| Manic Symptoms - Cho | 13 | rho = .654 | .015 |
| Negative Symptoms - mI | 7 | rho = -.771 | .042 |

**Table S8: Significant differences (p** ≤ **.05) in correlation coefficients: Schizophrenia (Scz) vs. Affective Psychosis (aff-P)**

| **Variables correlated** | **Scz correlation coefficient (rho)** | **Scz sample size** | **aff-P correlation coefficient (rho)** | **aff-P sample size** | **Z Statistic** | **P value (two-tailed)** |
| --- | --- | --- | --- | --- | --- | --- |
| Positive Symptoms - Cho | rho = -.523 | 20 | rho = .339 | 13 | -2.34 | .019 |
| Manic Symptoms - Cho | rho = -.114 | 20 | rho = .654 | 13 | -2.25 | .024 |
| Activation - Cho | rho = -.449 | 20 | rho = .352 | 13 | -2.14 | .032 |
| Autistic Preoccupation - Cho | rho = -.450 | 20 | rho = .317 | 13 | -2.04 | .041 |

**Table S9: Significant differences (p** ≤ **.05) in correlation coefficients: Healthy Controls (HC) vs. Schizophrenia (Scz)**

| **Variables correlated** | **HC correlation coefficient (rho)** | **HC sample size** | **Scz correlation coefficient (rho)** | **Scz sample size** | **Z Statistic** | **P value (two-tailed)** |
| --- | --- | --- | --- | --- | --- | --- |
| mI - depression | rho = .549 | 20 | rho = -.250 | 14 | 2.25 | .024 |
| NAA - activation | rho = .184 | 25 | rho = -.540 | 20 | 2.45 | .014 |
| Cho - activation | rho = .226 | 25 | rho = -.449 | 20 | 2.20 | .027 |
| mI - activation | rho = .319 | 20 | rho = -.842 | 14 | 4.03 | .0001 |
| Cr - dysphoria | rho = .070 | 25 | rho = -.607 | 20 | 2.40 | .016 |
| Cho - dysphoria | rho = .172 | 25 | rho = -.447 | 20 | 2.03 | .042 |

**Table S10: Significant differences (p** ≤ **.05) in correlation coefficients: Healthy Controls (HC) vs. Affective Psychosis (aff-P)**

| **Variables correlated** | **HC correlation coefficient (rho)** | **HC sample size** | **Scz correlation coefficient (rho)** | **Scz sample size** | **Z Statistic** | **P value (two-tailed)** |
| --- | --- | --- | --- | --- | --- | --- |
| Cho – manic symptoms | rho = -.053 | 26 | rho = .654 | 13 | -.221 | .027 |

**Table S11: Significant differences (p** ≤ **.05) in correlation coefficients: Non-psychotic Affective Disorders (NP-aff) vs. Schizophrenia (Scz)**

| **Variables correlated** | **NP-aff correlation coefficient (rho)** | **NP-aff sample size** | **Scz correlation coefficient (rho)** | **Scz sample size** | **Z Statistic** | **P value (two-tailed)** |
| --- | --- | --- | --- | --- | --- | --- |
| mI - activation | rho = -.176 | 16 | rho = -.842 | 14 | 2.56 | .011 |
| NAA - depression | rho = .308 | 22 | rho = -.449 | 22 | 2.40 | .016 |

**Table S12: Significant differences (p** ≤ **.05) in correlation coefficients: Non-psychotic Affective Disorders (NP-aff) vs. Affective Psychosis (aff-P)**

| **Variables correlated** | **NP-aff correlation coefficient (rho)** | **NP-aff sample size** | **aff-P correlation coefficient (rho)** | **aff-P sample size** | **Z statistic** | **P value (two-tailed)** |
| --- | --- | --- | --- | --- | --- | --- |
| Cho – manic symptoms | rho = -.028 | 22 | rho = .654 | 13 | -.207 | .039 |

**Table S13: Intercorrelations of neurometabolites in Schizophrenia (Scz)**

| **Variables correlated** | **Sample size** | **Correlation coefficient (rho)** | **P value (two-tailed)** |
| --- | --- | --- | --- |
| NAA - Cho | 20 | rho = .699 | <.001 |
| NAA - mI | 14 | rho = .762 | .002 |
| NAA- Glx | 10 | rho = .709 | .022 |
| NAA - Cr | 20 | rho = .430 | .058 |
| Cho - mI | 14 | rho = .642 | .013 |
| Cho - Glx | 10 | rho = .622 | .055 |
| Cho - Cr | 20 | rho = .615 | .004 |
| mI - Glx | 9 | rho = .921 | <.001 |
| mI - Cr | 14 | rho = .809 | <.001 |
| Glx - Cr | 10 | rho = .644 | .044 |
